# Loss of Endothelial HIF-Prolyl hydroxylase 2 (PHD2) Induces Cardiac Hypertrophy and Fibrosis

**DOI:** 10.1101/2021.03.19.434301

**Authors:** Zhiyu Dai, Jianding Cheng, Bin Liu, Dan Yi, Anlin Feng, Ting Wang, Chen Gao, Yibin Wang, Maggie M. Zhu, Xianming Zhang, You-Yang Zhao

## Abstract

**Background:** Cardiac hypertrophy and fibrosis are common adaptive responses to injury and stress, eventually leading to heart failure. Hypoxia signaling is important to the (patho)physiological process of cardiac remodeling. However, the role of endothelial Prolyl-4 hydroxylase 2 (PHD2)/hypoxia inducible factors (HIFs) signaling in the pathogenesis of heart failure remains elusive.

**Methods and Results:** Mice with *Tie2*-Cre-mediated deletion of *Egln1* (encoding PHD2) (*Egln1*^*Tie2Cre*^) exhibited left ventricular (LV) hypertrophy evident by increased thickness of anterior and posterior wall and LV mass, as well as cardiac fibrosis. Tamoxifen-induced endothelial *Egln1* deletion in adult mice also induced LV hypertrophy and fibrosis. Additionally, we observed a marked decrease of PHD2 expression in heart tissues and cardiovascular endothelial cells from patients with cardiomyopathy. Moreover, genetic ablation of *Hif2a* but not *Hif1a* in *Egln1*^*Tie2Cre*^ mice normalized cardiac size and function. RNA sequencing analysis also demonstrated HIF-2α as a critical mediator of signaling related to cardiac hypertrophy and fibrosis. Pharmacological inhibition of HIF-2α attenuated cardiac hypertrophy and fibrosis in *Egln1*^*Tie2Cre*^ mice.

**Conclusions:** The present studies define for the first time an unexpected role of endothelial PHD2 deficiency in inducing cardiac hypertrophy and fibrosis in a HIF-2α dependent manner. PHD2 was markedly decreased in cardiovascular endothelial cells in patients with cardiomyopathy. Thus, targeting PHD2/HIF-2α signaling may represent a novel therapeutic approach for the treatment of pathological cardiac hypertrophy and failure.

## Introduction

During development, the heart grows by cardiomyocyte proliferation and hypertrophy while after birth, the cardiomyocytes lose proliferative potential and heart growth is mainly via cardiomyocyte hypertrophy. Cardiac hypertrophy can happen in physiological and pathophysiological conditions. Pathological hypertrophy induced by hypertension, myocardial infarction, cardiomyopathy results in ventricular remodeling which is associated with systolic and diastolic dysfunction and interstitial fibrosis, and finally leads to deleterious outcomes such as heart failure^1,2^. Understanding the mechanistic molecular signaling in the event of physiological and pathological cardiac hypertrophy will lead to identify novel therapeutic approaches for patients with heart failure. There are multiple cells types including cardiomyocytes, endothelial cells (ECs), fibroblasts and smooth muscle cells in the heart. ECs lining of the inner layer of the blood vessels account for the greatest number of cells in the heart. One of the major functions of, cardiac vascular ECs is to supply oxygen and nutrients to support cardiomyocytes^3,4^. Previous studies have demonstrated that angiogenesis stimulated by VEGF-B or PIGF induces marked increase of cardiac mass in rodents, whereas inhibition of angiogenesis results in decreased capillary density, contractile dysfunction, and impaired cardiac growth^5,6^. Cardiac vasculature rarefaction is associated with pathological cardiac hypertrophy and heart failure^7^. Recent studies have also demonstrated the important role of EC-derived paracrine factors such as Endothelin-1, Apelin, neuregulin and Agrin in regulating cardiac hypertrophy, regeneration, and repair^8,9^. However, how ECs crosstalk with cardiomyocytes in the pathogenesis of cardiac hypertrophy and dysfunction is not fully understood.

Hypoxia-inducible factors (HIF) are key transcriptional factors mediating cellular response to oxygen levels. The α-subunit of HIF is delicately controlled by HIF-prolyl hydroxylases (PHD1-3)^10–12^. Under normoxia condition, PHDs hydroxylate HIF-α (mainly HIF-1α and HIF-2α), then E3 ligase VHL promotes degradation of hydroxylated HIF-α through proteasome-degradation pathway. Under hypoxia condition, PHD activities are inhibited leading to stabilization of HIF-α proteins which activate downstream target gene expression. Both HIF-1a and HIF-2α share similar DNA binding site or hypoxia response element (HRE)^11,13^. Thus, some genes are co-regulated by HIF-1α and HIF-2α, such as CXCL12. However, HIF-1α and HIF-2α also control some sets of unique downstream targets, for example, LDHA, is a HIF-1α target while OCT4 is a HIF-2α target^14^. HIF-1α is ubiquitously expressed whereas HIF-2α is more restricted in certain cell types such as ECs, and alveolar type 2 epithelial cells^15^, suggesting that HIF-α have distinct function in different cells under different (patho)physiological conditions. Our previous studies have shown that HIF-2α but not HIF-1α activation causes severe pulmonary hypertension in *Egln1* (encoding PHD2) conditional knockout mice while HIF-1α induction and activation is important for endothelial regeneration and vascular repair following inflammatory injury.^16,17^

*Egln1* null mutant mice exhibit polycythemia and congestive heart failure^18^. However, cardiomyocyte specific-disruption of *Egln1* causes only mild abnormality with the presence of occasional myocytes with increased hypereosinophilia and blurring of the cross-striations^19^, indicating loss of PHD2 cells other than cardiomyocytes causes congestive heart failure. Fan et al. show that EC deletion of both PHD2 and 3 induces spontaneous cardiomegaly due to enhanced cardiomyocyte proliferation and also prevents heart failure induced by myocardial infarction^20^. However, it is unknown whether these phenotypes are mediated by endothelial loss of either PHD2 or PHD3 alone. Studies have shown loss of both PHD2 and 3 but not PHD2 alone in cardiomyocyte induces ischemic cardiomyopathy^20^. Thus, it is important to determine whether loss of PHD2 alone in ECs affects cardiomyocyte and heart function in adult mice. To our surprise, we observed spontaneous left ventricular (LV) hypertrophy and cardiac fibrosis in tamoxifen-inducible EC-specific *Egln1* knockout adult mice. Using the mice with endothelial deletion of *Egln1* alone, *Egln1* and *Hif1a*, or *Egln1* and *Hif2a*, we found that loss of endothelial PHD2-induced cardiac hypertrophy and fibrosis was mediated by HIF-2α activation and pharmacological inhibition of HIF-2α inhibited LV hypertrophy and cardiac fibrosis. Analysis of single cell RNA sequencing datasets revealed *EGLN1* mainly expressed in vascular ECs of the human heart under normal condition. In patients with hypertrophic cardiomyopathy, *EGLN1* expression in LV heart tissue (mRNA) and cardiac vascular ECs (protein) was markedly deceased, which validate the clinical relevance of our findings that endothelial PHD2 deficiency induces LV hypertrophy and cardiac fibrosis. Thus, targeting PHD2/HIF-2α signaling could be a therapeutic approach for pathological cardiac hypertrophy leading to cardiomyopathy and heart failure.

## Methods

### Human Samples and Data

Archived human heart sections from the National Center for Medico-legal Expertise of Sun Yat-sen University were used. The patient heart samples were collected from six cardiomyopathy patients including four hypertrophic cardiomyopathy, one dilated cardiomyopathy, and one hypertensive cardiomyopathy. Control samples were collected from age- and gender-matched individuals (six males, one female; 27-53 years old) without heart disease. The cause of death for controls was mechanical injuries. All data and sample collection were approved by the ethics committee of Sun Yat-sen University. Informed consent was obtained from the legal representatives of the patients. *EGLN1* mRNA expression in heart tissues from patients with dilated cardiomyopathy were provided by Drs. Chen Gao and Yibin Wang^21,22^. Failing heart samples were obtained from the left ventricular anterior wall during heart transplantation or implantation of LVAD (LV assistant device). The non-failing heart samples were obtained from the LV free wall and procured from National Disease Research Interchange and University of Pennsylvania. Non-failing heart donors showed no laboratory signs of cardiac disease. Tissue collection was approved by UCLA Institutional Review Board #11-001053; #12-000207.

### Animals

*Egln1*^*Tie2Cre*^ mice, *Egln1*^*Tie2Cre*^*/Hif1a*^*Tie2Cre*^ and *Egln1*^*Tie2Cre*^*/Hif2a*^*Tie2Cre*^ double knock out mice were generated as described previously^16^. *Egln1*^*EndoCreERT2*^ mice were generated by crossing *Egln1* floxed mice with *EndoSCL-CreERT2* transgenic mice expressing the tamoxifen-inducible *Cre* recombinase under the control of the 5’ endothelial enhancer of the stem cell leukemia locus^23^. At 8 weeks of age, *Egln1*^*EndoCreERT2*^ and *Egln1*^*f/f*^ mice were treated with 2 mg tamoxifen intraperitoneally for 5 days to induced *Egln1* deletion only in ECs. Mice were scarified at the age of ∼9 months. Both male and female mice were used in these studies. The use of animals was in compliance with the guidelines of the Animal Care and Use Committee of the Northwestern University and the University of Arizona.

### Echocardiography

Echocardiography was performed on a Fujifilm VisualSonics Vevo 2100 using an MS550D (40 MHz) transducer as described previously. Briefly, mice were anesthetized in an induction chamber filled with 1% isoflurane. The left ventricle anterior/posterior wall thickness during diastole (LVAW and LVPW), the LV internal dimension during diastole (LVID), the LV fractional shortening (LV FS) and the cardiac output (CO) were obtained from the parasternal short axis view using M-mode. Results were calculated using VisualSonics Vevo 2100 analysis software (v. 1.6) with a cardiac measurements package and were based on the average of at least three cardiac cycles.

### Reanalysis of public single-cell RNA sequencing datasets

We used the publicly available metadata from two human fetal hearts (GSM4008686 and GSM4008687)^24^. The metadata was processed in R (version 4.0.2) via *Seurat* package V3.2.3^25^. Briefly, cells that expressed fewer than 100 genes, and cells that expressed over 4,000 genes, and cells with unique molecular identifiers (UMIs) more than 10% from the mitochondrial genome were filtered out. The data were normalized and integrated in *Seurat*, followed by Scaled and summarized by principal component analysis (PCA), and then visualized using UMAP plot. *FindClusters* function (resolution = 0.5) in Seurat was used to cluster cells based on the gene expression profile. Cardiomyocyte (*NPPA, TNNT2*), endothelial cells (*CDH5*), fibroblasts (*FN1, VIM*), smooth muscle cells (*ACTA2*), and macrophages (*CD68, CD14*) clusters were annotated based on the expression of known markers.

### Irradiation and Bone marrow transplantation

WT or *Egln1*^*Tie2Cre*^ female mice at the age of 3 weeks were delivered at a dose of 750 cGy/mouse. At 3 hours following irradiation, mice were transplanted with 1×10^7^ bone marrow cells (in 150 μl of HBSS) isolated from male *Egln1*^*Tie2Cre*^ or WT mice through tail vein injection. Mice were used for heart dissection at the age of 3.5 months as described previously^16^.

### Immunofluorescent, immunohistochemical and histological staining

Following PBS perfusion, heart tissue was embedded in OCT for cryosectioning for immunofluorescent staining. Heart sections (5 μm) were fixed with 4% paraformaldehyde, followed by blocking with 0.1% Triton X-100 and 5% normal goat serum at room temperature for 1 h. After 3 washes, they were incubated with anti-Ki67 (1:25, Abcam, Cat#ab1667), anti-CD31 antibody (1:25, BD Biosciences, Cat#550274) at 4 °C overnight and were then incubated with either Alexa 647-conjugated anti-rabbit IgG (Life Technology), or Alexa 594-conjugated anti-rabbit IgG or Alexa 488 conjugated anti-rat IgG at room temperature for 1 h after 3 washes. Nuclei were counterstained with DAPI contained in Prolong Gold mounting media (Life Technology).

For Wheat Germ Agglutinin (WGA) staining, WGA conjugated with FITC or WGA conjugated with Alexa 647 were stained with cryosectioned slides at room temple for 10 mins. For immunohistochemistry staining on paraffin sections, the tissues were cut into semi-thin 5 μm thick sections after paraffin processing. Heart sections were then dewaxed and dehydrated. Antigen retrieval was performed by boiling the slides in 10 mmol/L sodium citrate (pH 6.0) for 10 minutes. After blocking, slides were incubated with anti-PHD2 antibody (1:200, Cell Signaling Technology, Cat#4835), at 4 °C overnight. Slides were incubated with 6% H2O2 for 30 min after primary antibody incubation and were then biotinylated with a rabbit IgG and ABC kit (Vector Labs) for immunohistochemistry. The nucleus was co-stained with hematoxylin (Sigma-Aldrich).

For histological assessment, hearts were harvested and washed with PBS, followed by fixation in 4% formalin and dehydrated in 70% ethanol. After paraffin processing, the tissues were cut into semi-thin 5 m thick sections. Sections were stained with H & E staining or Masson’s trichrome kit as a service of charge at Core Facility.

### QRT-PCR

Total RNA was isolated from frozen left ventricular tissues with Trizol reagents (Invitrogen) followed by purification with the RNeasy Mini kit including DNase I digestion (Qiagen). One microgram of RNA was transcribed into cDNA using the high-capacity cDNA reverse transcription kits (Applied Biosystems) according to the manufacturer’s protocol. Quantitative RT-PCR analysis was performed on an ABI ViiA 7 Real-time PCR system (Applied Biosystems) with the FastStart SYBR Green Master kit (Roche Applied Science). Target mRNA was determined using the comparative cycle threshold method of relative quantitation. *Cyclophilin* was used as an internal control for analysis of expression of mouse genes. The primer sequences are provided in **Supplemental Table 1**.

### RNA-sequencing

Total RNA was isolated from left ventricular tissues with Trizol reagents (Invitrogen) followed by purification with the RNeasy Mini kit including DNase I digestion (Qiagen). RNA sequencing was carried out by Novogene corporation on Illumina Hiseq platform. The original sequencing data were trimmed using FASTX and aligned to the reference genome using TopHat2. The differential expression analysis was performed using Cuffdiff software^26^.

### Data availability

Scripts used for single-cell RNA sequencing analysis, analyzed data in R objects, and the RNA sequencing raw data are available in Figshare (https://figshare.com/projects/Loss_of_Endothelial_HIF-Prolyl_hydroxylase_2_PHD2_Induces_Cardiac_Hypertrophy_and_Fibrosis/98861).

### Statistics

Statistical analysis of data was done with Prism 7 (GraphPad Software, Inc.). Statistical significance for multiple-group comparisons was determined by one-way ANOVA with Tukey post hoc analysis that calculates corrected P values. Two-group comparisons were analyzed by the unpaired two-tailed Student’s *t* test for equal variance or by Mann-Whitney test for unequal variance. *P* <0.05 denoted the presence of a statistically significant difference. All bar graphs represent means ±SD. Bars in dot plot figures represent the mean.

## Results

### Decreased PHD2 expression in patient with cardiomyopathy

PHDs/VHL/HIF signaling have been implicated in many physiological and pathological conditions of heart development and heart diseases. Leveraging the public single-cell RNA-sequencing dataset, we first analyzed the mRNA expression of the key molecules of PHDs/VHL/HIF signaling from fetal and adult hearts. Our data demonstrated that *EGLN1* and *EPAS1* (encoding HIF-2α) are highly expressed in cardiac ECs in both the fetal and adult hearts (**Figure 1A** and **supplemental Figure 1**). We then examined the expression levels of PHD2 in heart tissues of cardiomyopathy patients and normal donors by quantitative (Q)RT-PCR. *EGLN1* mRNA levels were drastically decreased in the LV cardiac tissue from patients with cardiomyopathy compared to normal donors (**Figure 1B**). To further determine the cell-specific loss of PHD2 expression in the heart, we performed immunohistochemistry on LV heart sections. PHD2 was highly expressed in ECs as well as smooth muscle cells but only mildly in cardiomyocytes in normal donor hearts. However, its levels were dramatically reduced in cardiovascular ECs but not in smooth muscle cells of cardiomyopathy patients (**Figure 1C** and **1D**). These data demonstrate a marked loss of endothelial PHD2 expression in patients with cardiomyopathy, suggesting a crucial role of endothelial PHD2 in cardiac function.

**Figure 1.**
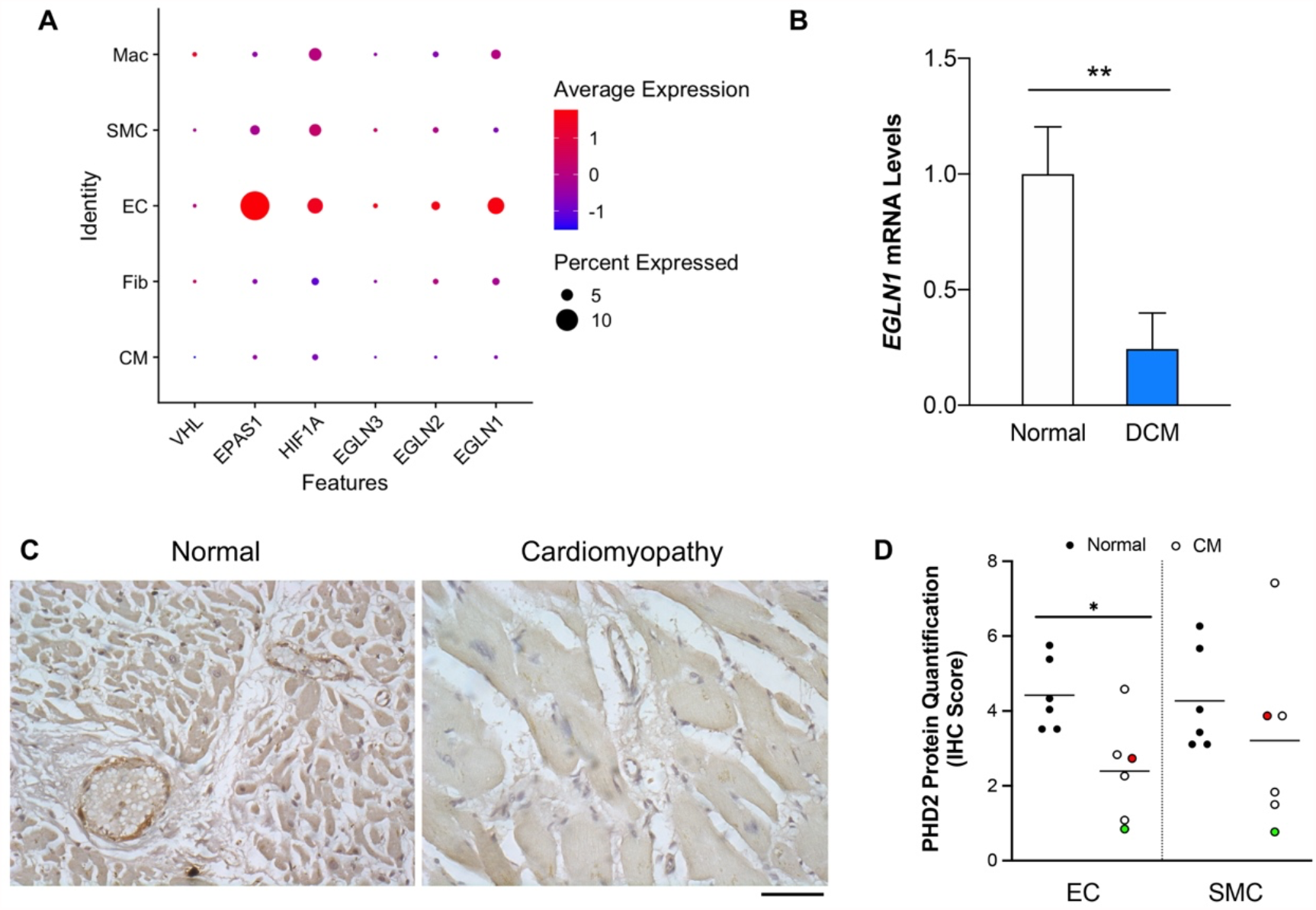
Decreased PHD2 expression in heart vascular ECs of patients with cardiomyopathy. (**A**) Single-cell RNA sequencing analysis from human fetal hearts showed that cardiac ECs express higher levels of *EGLN1, EPAS1* compared to other cell types. Mac=macrophage, SMC= smooth muscle cell, Fib=fibroblast, CM=cardiomyocyte. (**B**) QRT-PCR analysis showing that *EGLN1* mRNA levels in LV hearts were significantly downregulated in dilated cardiomyopathy (DCM) patients compared to healthy donors. N=3 each group. (**C** and **D**) Immunohistochemistry staining against PHD2 (**C**) and quantification (**D**) demonstrating a significant decrease of PHD2 expression in LV sections, especially in vascular endothelial cells (ECs) of patients with cardiomyopathy (CM). SMC indicates smooth muscle cell. White dots indicate hypertrophic cardiomyopathy, red dot indicate dilated cardiomyopathy, green dot indicates hypertensive cardiomyopathy. Scale bar, 50μm. \**p* < 0.05, \*\**p* <0.01 (Student’s *t* test). Immunostaining intensity was graded from 1 to 10, with 10 being the highest.

### Constitutive loss of endothelial PHD2 induces left ventricular hypertrophy and fibrosis

To investigate the role of endothelial PHD2 in the heart *in vivo*, we generated *Egln1*^*Tie2Cre*^ mice by breeding *Egln1* floxed mice with *Tie2Cre* transgenic mice. Dissection of cardiac tissue showed marked increase of the ratio of LV vs body weight (LV/BW), indicative of LV hypertrophy (**Figure 2A**). Echocardiography revealed marked increases of LV anterior and posterior wall thicknesses (**Figure 2B-2E**). There was no significant change of LV systolic function by evaluation of fractional shortening and ejection fraction (**Supplemental Figure 2A** and **2B**). Histological examination demonstrated that *Egln1*^*Tie2Cre*^ mice exhibited marked increase of wall thickness of the LV as well as the RV and reduction of the chamber sizes, indicating cardiac hypertrophy (**Figure 2F**). Wheat germ agglutinin (WGA) staining showed that LV cardiomyocytes from *Egln1*^*Tie2Cre*^ mice had marked increase of cellular surface, indicative of cardiomyocyte hypertrophy (**Figure 2G**). QRT-PCR analysis revealed a marked increase of expression of *Anp, Bnp* and *Myh7*, further supporting cardiac hypertrophy (**Supplemental Figure 3**). We also observed significant perivascular and inter-cardiomyocyte fibrosis in the LV of *Egln1*^*Tie2Cre*^ mice by Trichrome staining (**Figure 2H**). Consistently *Col1a* expression was also markedly increased in the LV of *Egln1*^*Tie2Cre*^ mice (**Supplemental Figure 3**).

**Figure 2.**
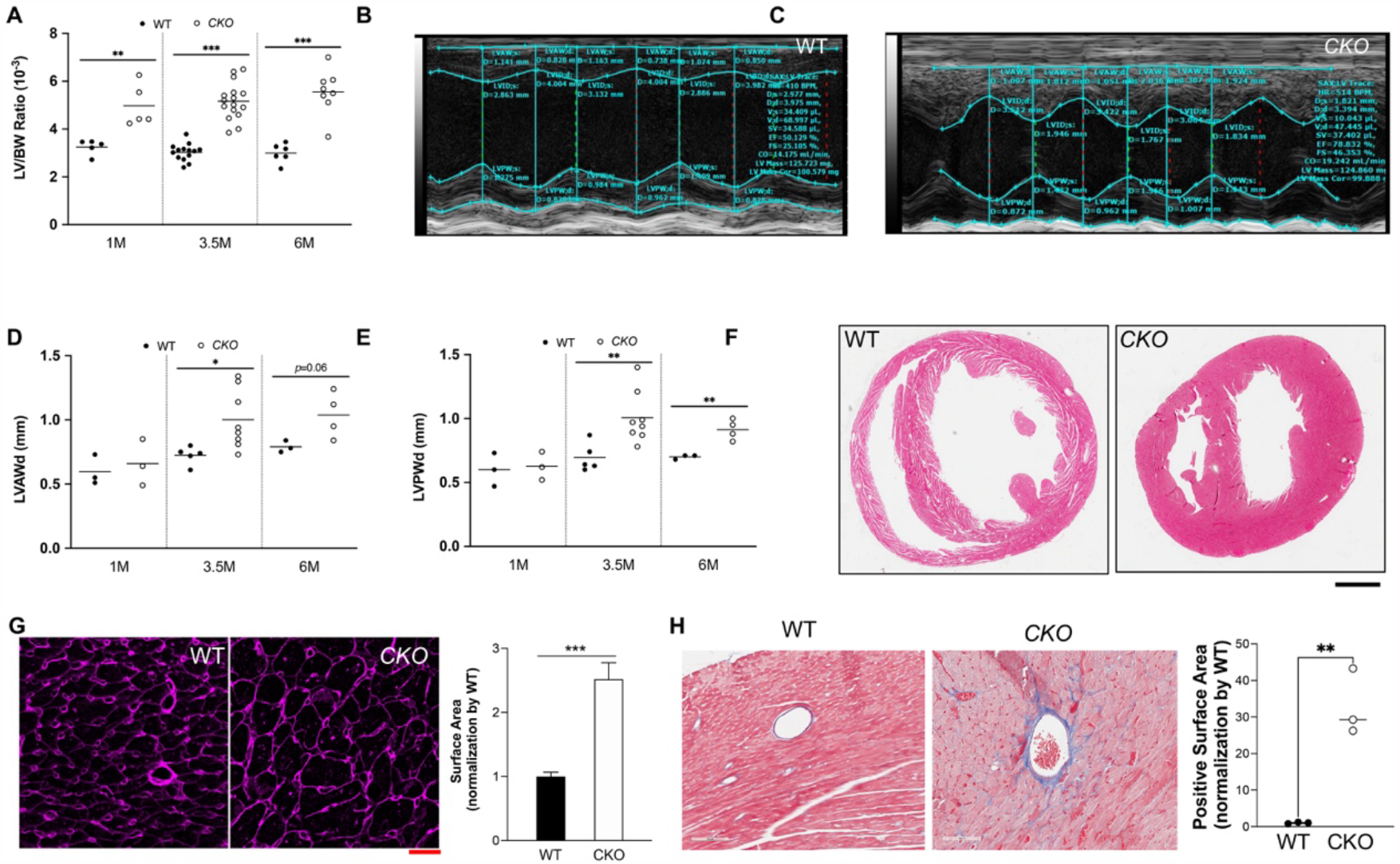
Constitutive loss of endothelial *Egln1* leads to left ventricular hypertrophy and fibrosis. (**A**) Increased weight ratio of LV/body weight of *Egln1*^*Tie2Cre*^ (CKO) mice compared to WT mice at various ages. M = month. (**B** and **C**) Representative M model of echocardiography measurement showing that *Egln1*^*Tie2Cre*^ mice exhibit increase of LV anterior, posterior wall thicknesses, and reduction of LV internal dimension (LVID) compared to WT mice at the age of 3.5 months. (**D** and **E**) Quantification of LV anterior and posterior wall thicknesses via echocardiography measurement WT and *Egln1*^*Tie2Cre*^ mice. (**F**) H&E staining showing cardiac hypertrophy in *Egln1*^*Tie2Cre*^ mice at the age of 3.5 months. Scale bar, 1 mm. (**G**) WGA staining of cardiac sections and quantification demonstrating enlargement of cardiomyocytes in *Egln1*^*Tie2Cre*^ mice. N=5/group. Scale bar, 20μm. (**H**) Trichrome staining showing prominent cardiac fibrosis in *Egln1*^*Tie2Cre*^ mice. Scale bar, 100μm. * *p* <0.05, ** *p* <0.01, *** *p* <0.001 (Student’s *t* test).

As *Tie2Cre* also expresses in hematopoietic cells in addition to ECs, we next determined whether hematopoietic cell-expressed PHD2 plays a role in inducing LV hypertrophy. We performed bone marrow transplantation via transplanting WT or *Egln1*^*Tie2Cre*^ bone marrow cells to lethally irradiated WT or *Egln1*^*Tie2Cre*^ mice. Three months post-transplantation, we did not observe any change of LV/BW ratio in WT mice transplanted with *Egln1*^*Tie2Cre*^ bone marrow cells compared to WT mice with WT bone marrow cells, indicating loss of PHD2 in bone marrow cells *per se* didn’t induce LV hypertrophy. Similarly, WT bone marrow cell transplantation to *Egln1*^*Tie2Cre*^ mice didn’t affect LV hypertrophy seen in *Egln1*^*Tie2Cre*^ mice transplanted with *Egln1*^*Tie2Cre*^ bone marrow cells (**Supplemental Figure 4A** and **4B**). These data suggest that loss of hematopoietic cell PHD2 is not involved in the development of LV hypertrophy.

### Inducible deletion of endothelial *Egln1* in adult mice leads to development of LV hypertrophy and fibrosis

To determine if the cardiac hypertrophy in the *Egln1*^*Tie2Cre*^ mice is ascribed to potential developmental defects in mice with *Tie2*Cre-mediated deletion of *Egln1*,we generated mice with tamoxifen-inducible endothelial *Egln1* deletion via breeding *Egln1* floxed mice with *EndoSCL-CreERT2* mice^27^ (**Figure 3A**), which induces EC-restricted gene disruption^23,28–30^. Tamoxifen was administrated to *Egln1*^*EndoCreERT2*^ mice at age of 8 weeks. Seven months post-tamoxifen treatment, echocardiography, cardiac dissection, and histological analysis were employed to evaluate the cardiac phenotype. *Egln1*^*EndoCreERT2*^ mice exhibited a marked increase of LV/BW ratio (**Figure 3B**), LV wall thicknesses and LV mass (**Figure 3C-3E**). Histologic analysis also demonstrated marked increase of LV wall thickness, reduction of LV chamber size, and perivascular fibrosis by Trichrome staining (**Figure 3F-3G**). However, the RV wall thickness and chamber size were not affected. Different from the *Egln1*^*Tie2Cre*^ mice by *Tie2*Cre-mediated deletion which also exhibit severe pulmonary hypertension^16,31,32^, the *Egln1*^*EndoCreERT2*^ mice had only a mild increase of right ventricular systolic pressure (data not shown) indicative mild pulmonary hypertension which is consistent with minimal changes in RV wall thickness and chamber size. These data support the idea that loss of endothelial PHD2 in adult mice selectively induces LV hypertrophy.

**Figure 3.**
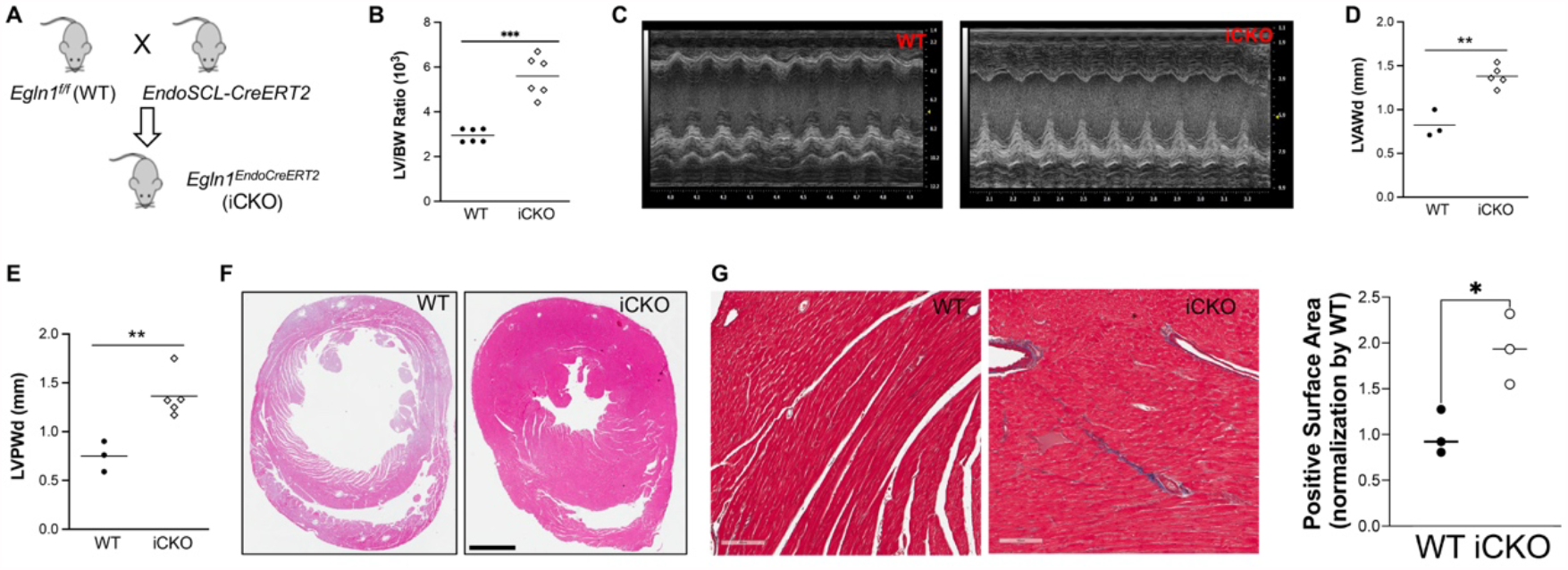
Inducible deletion of endothelial *Egln1* in adult mice leads to left ventricular hypertrophy and fibrosis. (**A**) A diagram showing the strategy of generating *Egln1*^*EndoCreERT2*^ mice (iCKO). (**B**) *Egln1*^*EndoCreERT2*^ mice exhibited increase of LV/body weight ratio compared to age-matched WT mice at ∼7 months post-tamoxifen treatment. (**C**) Representative M model of echocardiography showing LV hypertrophy in *Egln1*^*EndoCreERT2*^ mice. (**D** and **E**) Quantification of LVAWd and LVPWd showing increased LV wall thicknesses of *Egln1*^*EndoCreERT2*^ mice. (**F**) H&E staining demonstrating that *Egln1*^*EndoCreERT2*^ mice developed LV hypertrophy. Scale bar, 1mm. (**G**) Trichrome staining showing the deposition of collagen in the LV of *Egln1*^*EndoCreERT2*^ mice. Scale bar, 100μm.** *p* <0.01, *** *p* <0.001 (Student’s *t* test).

### Increased endothelial proliferation and angiogenesis in the LV of endothelial PHD2*-* deficient mice

Previous studies demonstrated that stimulation of angiogenesis in the absence of other insults can drive myocardial hypertrophy in mice via overexpression of PR39 or VEGF-B^5^. PHD2 deficiency has been shown to induce EC proliferation and angiogenesis *in vitro* and *in vivo*^33,34^. To determine if vascular mass is increased in the LV of the *Egln1*^*Tie2Cre*^ mice, we performed immunostaining with endothelial marker CD31 and WGA. The capillary/myocyte ratio was drastically increased in the LV of *Egln1*^*Tie2Cre*^ mice compared to WT mice (**Figure 4A** and **4B**). Anti-Ki67 immunostaining demonstrated a marked increase of Ki67^+^/CD31^+^ cells in the LV of *Egln1*^*Tie2Cre*^ mice (**Figure 4C** and **4D**), suggesting that PHD2 deficiency induces EC proliferation in the LV, which explains the marked increase of capillary/myocyte ratio.

**Figure 4.**
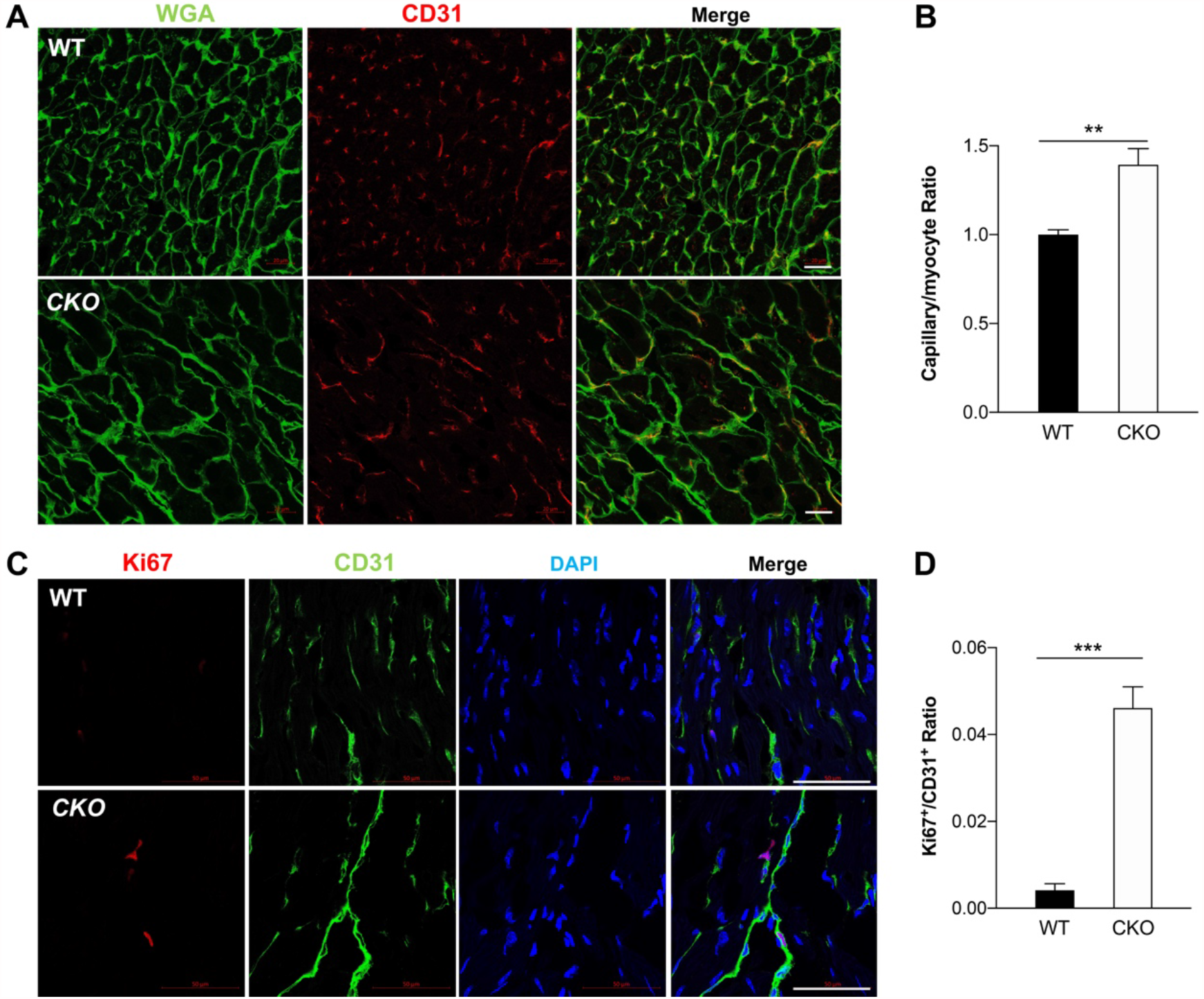
Deletion of endothelial *Egln1* increased angiogenesis in the left ventricles. (**A** and **B)** Immunostaining of CD31 and WGA and quantification showing increase of capillary EC versus cardiomyocyte number in *Egln1*^*Tie2Cre*^ mice. Left heart sections were stained with anti-CD31 (red, marker for ECs) and WGA (green). N=4/group. Scale bar, 20μm. (**C** and **D**) Anti-Ki67 staining, and quantification demonstrated that cardiac EC was hyperproliferative in the LV of *Egln1*^*Tie2Cre*^ mice. Left heart sections were immunostained with anti-Ki67 (red, cell proliferation marker) and anti-CD31 (green). Nuclei were counterstained with DAPI. N=4/group. Scale bar, 50μm. ** *p* <0.01, *** *p* <0.001 (Student’s *t* test).

### Distinct role of endothelial HIF-1α versus HIF-2α in LV hypertrophy

As *Egln1* deletion stabilizes both HIF-1α and HIF-2α, we generated *Egln1/Hif1a*^*Tie2Cre*^, *Egln1/Hif2a*^*Tie2Cre*^ double knock out and *Egln1/Hif1a/Hif2a*^*Tie2Cre*^ triple knock out mice (**Figure 5A**) to determine the HIF-α isoform(s) mediating LV hypertrophy. Heart dissection showed that *Egln1/Hif2a*^*Tie2Cre*^ and *Egln1/Hif1a/Hif2a*^*Tie2Cre*^ mice exhibited normal LV/BW ratio seen in WT mcie, whereas *Egln1/Hif1a*^*Tie2Cre*^ mice had increased LV/BW ratio compared to *Egln1*^*Tie2Cre*^ mice (**Figure 5B**). These data provide *in vivo* evidence that endothelial HIF-2α activation secondary to loss of endothelial PHD2 is responsible for LV hypertrophy seen in *Egln1*^*Tie2Cre*^ mice whereas HIF-1α activation attenuates LV hypertrophy. Echocardiography confirmed that *Egln1/Hif2a*^*Tie2Cre*^ mice had normal LV wall thickness and LV mass in contrast to *Egln1*^*Tie2Cre*^ mice (**Figure 5C** and **5D**). Histological assessment demonstrated that *Egln1/Hif2a*^*Tie2Cre*^ mice had no cardiac hypertrophy and fibrosis, as well as normal cardiomyocyte surface area (**Figure 5E**-**5G**). These data demonstrate the causal role of endothelial HIF-2α activation in mediating LV hypertrophy and fibrosis seen in *Egln1*^*Tie2Cre*^ mice.

**Figure 5.**
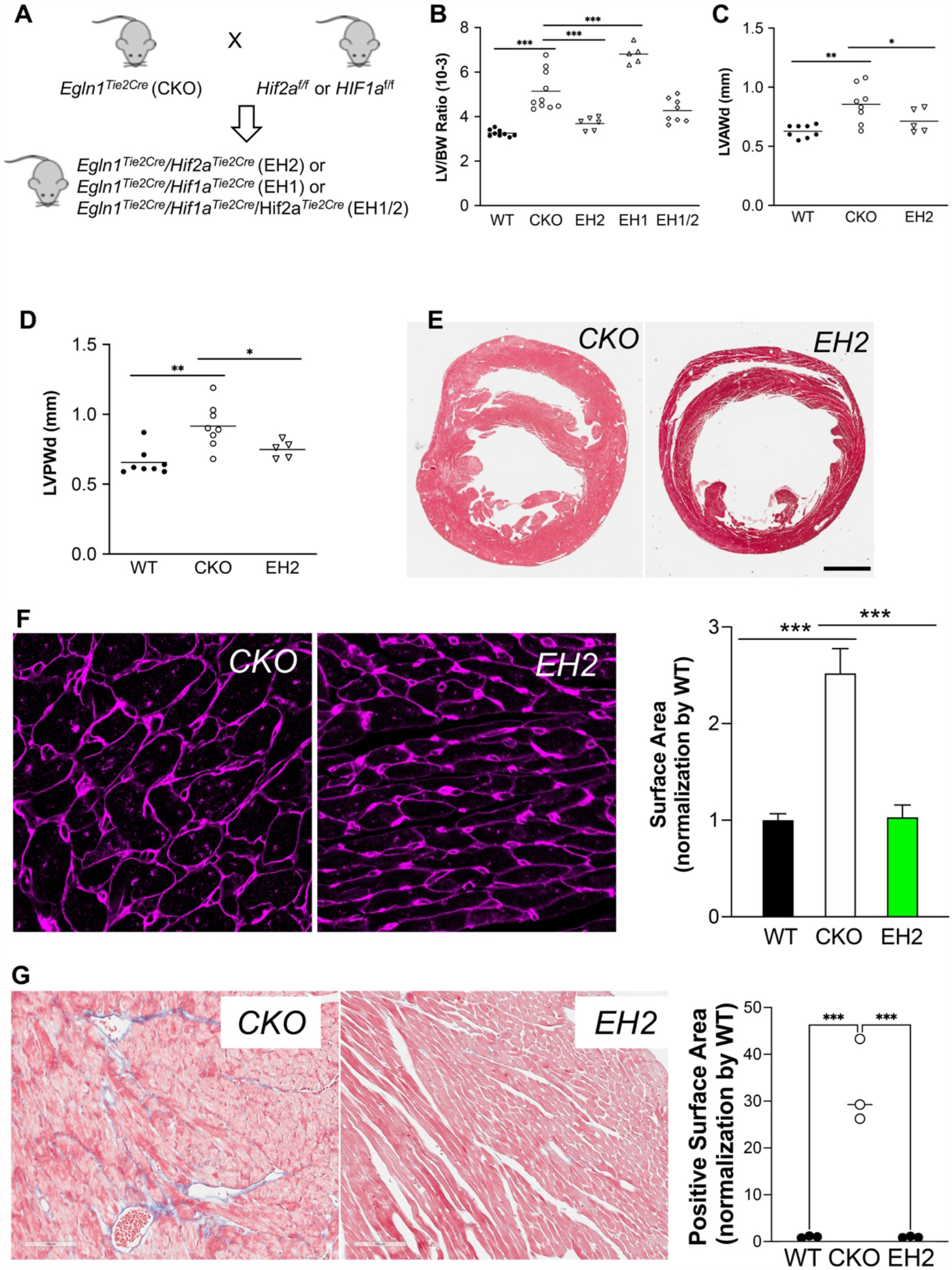
Distinct role of endothelial HIF-1α and HIF-2α in *Egln1* deficiency-induced left heart hypertrophy and fibrosis. **(A)** A diagram demonstrating generation of *Egln1/Hif2a*^*Tie2Cre*^ (EH2), *Egln1/Hif1a*^*Tie2Cre*^ (EH1) and *Egln1/Hif1a*/*Hif2a*^*Tie2Cre*^ (EH1/2) mice. (**B**) Cardiac dissection showing that endothelial HIF-2α deletion protected from endothelial *Egln1* deficiency-induced LV hypertrophy, whereas HIF-1α deletion augmented LV hypertrophy. (**C** and **D**) Echocardiography demonstrated that HIF-2α deletion in ECs protected from LV wall thickening induced by *Egln1* deficiency. (**E**) H&E staining showed normalization of LV hypertrophy in *Egln1/Hif2a*^*Tie2Cre*^ mice. Scale bar, 1mm. (**F**) WGA staining and quantification showed a complete normalization of cardiomyocyte hypertrophy in *Egln1/Hif2a*^*Tie2Cre*^ mice. The same surface area data of WT and KO in Fig.2H was used. N=5/group. Scale bar, 20μm. (**G**) Trichrome staining demonstrated absence of collagen deposition in the LV of *Egln1/Hif2a*^*Tie2Cre*^ mice. Scale bar, 100μm. * *p* <0.05, ** *p* <0.01, *** *p* <0.001 (One-way *ANOVA* with Tukey post hoc analysis for multiple group comparisons).

### RNA-sequencing analysis identifies HIF-2α-mediated signaling pathways regulating cardiac hypertrophy and fibrosis

To understand the downstream mechanisms of endothelial HIF-2α in regulating cardiac function, we performed whole transcriptome RNA sequencing (RNA-seq) of LV tissue dissected from WT, *Egln1*^*Tie2Cre*^ and *Egln1/Hif2a*^*Tie2Cre*^ mice (**Figure 6A**). Firstly, we found 1,454 genes (FPKM>5, fold change >1.5 or <0.66) were changed in *Egln1*^*Tie2Cre*^ lungs compared to WT LV. Then, we performed the enriched Kyoto Encyclopedia of Genes and Genomes (KEGG) pathways analysis on the upregulated genes in *Egln1*^*Tie2Cre*^ vs WT mice. The analysis revealed that there were alterations of upregulated pathways related to adrenergic signaling in cardiomyocytes, hypertrophic cardiomyopathy, dilated cardiomyopathy, and cardiac muscle contraction, which are consistent with the hypertrophic phenotype we observed in *Egln1*^*Tie2Cre*^ mice (**Figure 6B**). To determine the genes downstream of HIF-2α activation, we did the intersecting analysis of DEGs from WT_ *Egln1*^*Tie2Cre*^ and *Egln1*^*Tie2Cre*^_ *Egln1/Hif2a*^*Tie2Cre*^. There were 864 overlapping genes between WT_ *Egln1*^*Tie2Cre*^ and *Egln1*^*Tie2Cre*^_ *Egln1/Hif2a*^*Tie2Cre*^. KEGG pathway analysis of these genes upregulated in *Egln1*^*Tie2Cre*^ but normalized in *Egln1/Hif2a*^*Tie2Cre*^ hearts also showed enrichment of hypertrophic cardiomyopathy, and dilated cardiomyopathy (**Figure 6C)**, suggesting that these pathway abnormalities are downstream of HIF-2α activation. We also observed that many genes related to endothelium and EC-derived factors-mediated cell growth, and genes related to cardiac fibrosis were altered in *Egln1*^*Tie2Cre*^ mice whereas normalized in *Egln1/Hif2a*^*Tie2Cre*^ (**Figure 6D**). These data provide mechanistic understanding of the distinct roles of endothelial HIF-1α and HIF-2α in regulating LV hypertrophy and fibrosis.

**Figure 6.**
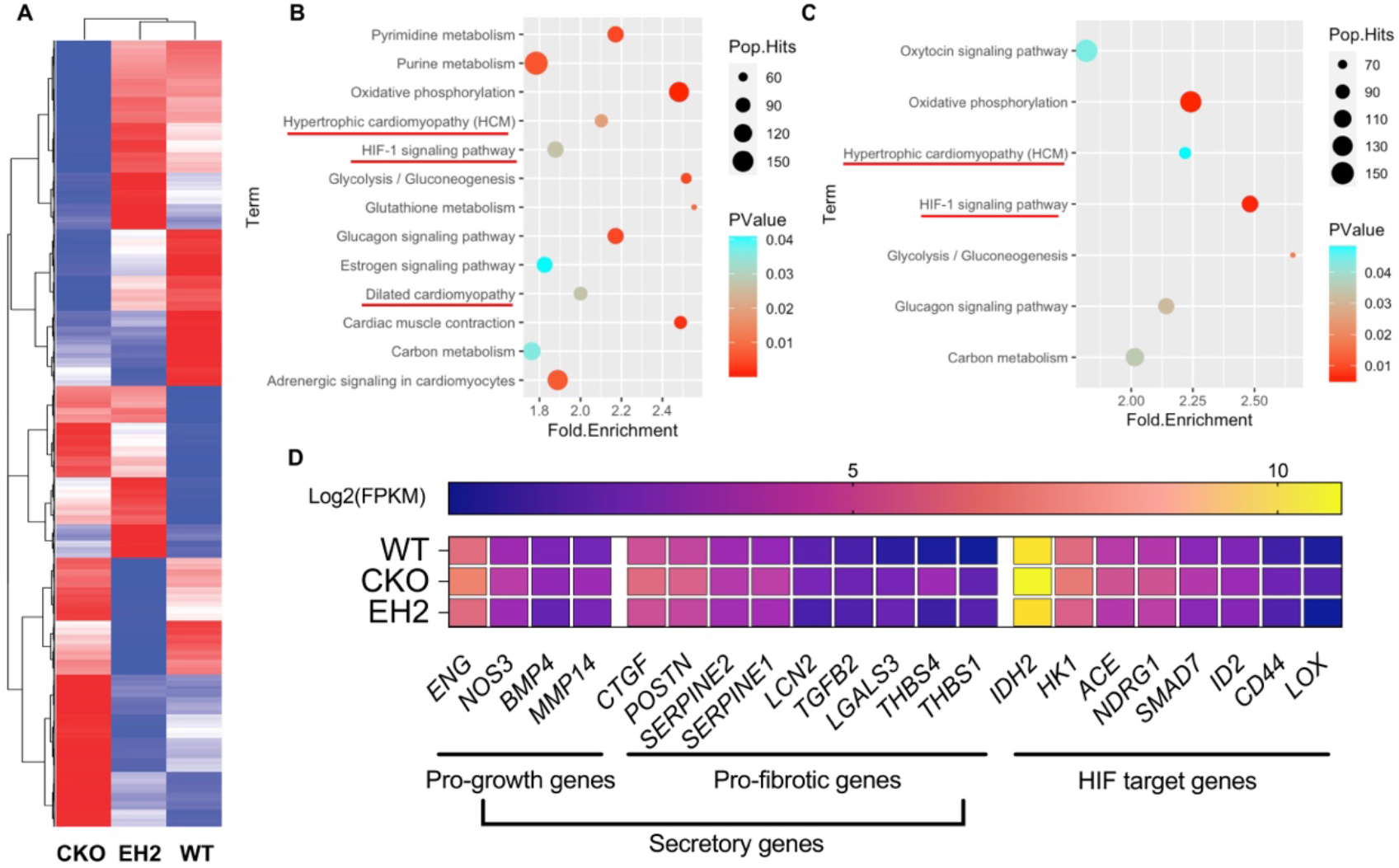
RNA-sequencing analysis identifies multiple dysfunctional pathways in regulation of cardiac hypertrophy and fibrosis induced by endothelial *Egln1* deficiency and normalization by HIF-2α disruption. **(A)** A representative heatmap of RNA-sequencing analysis of WT, *Egln1*^*Tie2Cre*^ (CKO) and *Egln1/Hif2a*^*Tie2Cre*^ (EH2) mice. (**B**) KEGG pathway enrichment analysis of upregulated genes in *Egln1*^*Tie2Cre*^ vs WT mice showing dysregulation of multiple signaling pathways related to cardiac hypertrophy. (**C**) Intersecting analysis of differential expression genes (DEG) between WT_*Egln1*^*Tie2Cre*^ and *Egln1*^*Tie2Cre*^_ *Egln1/Hif2a*^*Tie2Cre*^ mice. (**D**) KEGG pathway enrichment analysis indicates that HIF-2α-activated downstream genes are related to hypertrophic cardiomyopathy. (**E**) A heatmap showing the expression of genes related to endothelium and endothelial cell-derived factors-mediated cell growth, genes related to cardiac fibrosis, and HIF target genes.

### Pharmacological inhibition of HIF-2α inhibits LV hypertrophy

To assess the therapeutic potential of HIF-2α inhibition for cardiomyopathy, we treated 3-week-old *Egln1*^*Tie2Cre*^ mice with the HIF-2α translation inhibitor C76^31,35^ for 11 weeks. C76 treatment inhibited LV hypertrophy of *Egln1*^*Tie2Cre*^ mice evident by decreased LV/BW ratio compared to vehicle treatment (**Figure 7A**). C76 treatment also significantly attenuated the cardiomyocyte surface area assessed by WGA staining (**Figure 7B** and **7C**), and perivascular fibrosis by trichrome staining (**Figure 7D**). Taken together, these data demonstrated that inhibition of HIF-2α via a pharmacologic approach suppressed endothelial PHD2 deficiency-induced LV hypertrophy and cardiac fibrosis.

**Figure 7.**
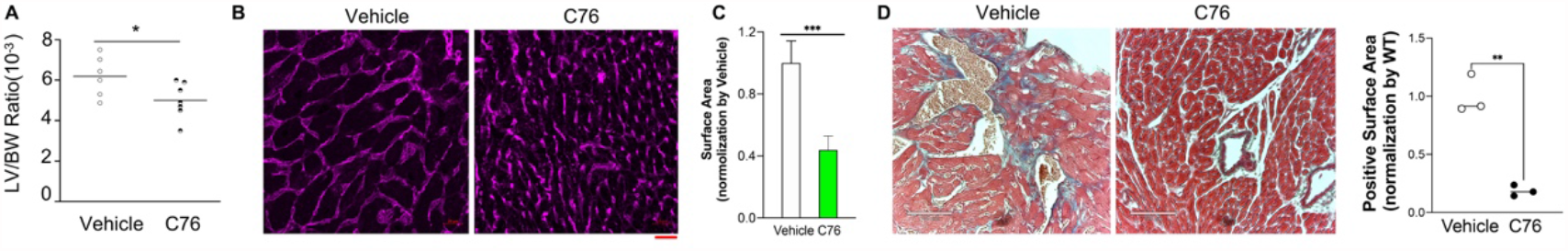
Pharmacological inhibition of HIF-2α attenuated cardiac hypertrophy and fibrosis. (A) Inhibition of HIF-2α reduced weight ratio of LV/body weight (BW) in *Egln1*^*Tie2Cre*^ mice. (**B** and **C**) WGA staining and quantification demonstrated that HIF-2α inhibition reduced cardiomyocyte hypertrophy. N=5/group. Scale bar, 20μm. (D) Trichrome staining revealed attenuation of cardiac fibrosis by HIF-2α inhibition. Scale bar, 100μm. * *p* <0.05, *** *p* <0.001 (Student’s *t* test).

## Discussion

In this study we demonstrate that endothelial PHD2 is markedly decreased in the hearts of patients with cardiomyopathy. Genetic deletion of endothelial *Egln1* in mice induces spontaneously severe cardiac hypertrophy and fibrosis. Through genetic deletion of *Hif2a or Hif1a* deletion in *Egln1*^*Tie2Cre*^ mice, we also demonstrate that endothelial HIF-2α but not HIF-1α activation is responsible for PHD2 deficiency-induced LV hypertrophy and fibrosis. Moreover, pharmacological inhibition of HIF-2α reduces cardiac hypertrophy in *Egln1*^*Tie2Cre*^ mice. Thus, our data provide strong evidence that endothelial homeostasis is crucial to maintain normal cardiac function.

We demonstrate that PHD2 is mainly expressed in the vascular endothelium in human heart under normal condition, and its expression is markedly reduced in cardiac tissue and ECs of patients with cardiomyopathy. So far as we know, our study for the first time indicates the clinical importance of PHD2 deficiency in the pathogenesis of cardiomyopathy and heart failure.

Mice with *Tie2*Cre-mediated deletion of *Egln1* develop LV hypertrophy and also RV hypertrophy associated with severe pulmonary hypertension^16,31^. The RV hypertrophy is secondary to marked increase of pulmonary artery pressure. Furthermore, we employed the tamoxifen-inducible *Egln1* knockout mice to determine the role of endothelial PHD2 deficiency in heart function. Seven months after tamoxifen treatment of 8 weeks old adult mice, we observed only LV hypertrophy in the mutant mice with quite normal RV size. These data provide clear evidence that endothelial PHD2 deficiency in adult mice selectively induces LV hypertrophy. Published studies have shown that loss of both endothelial PHD2 and PHD3 leads to enhanced ejection fraction and LV cardiomegaly due to increased cardiomyocyte proliferation^20^. Our study demonstrates that loss of endothelial PHD2 alone is sufficient to induce LV hypertrophy without marked changes in cardiomyocyte proliferation and LV contractility and ejection fraction. We also observed marked LV cardiac fibrosis with predominantly in the peri-vascular regions and less in inter-cardiomyocytes area, which may lead to heart dysfunction. Cardiac hypertrophy is associated with fibrosis indicates maladaptive deleterious remodeling^36^. However, our data did not suggest that fibrosis is associated with heart failure, which might be due to the variability among animals. For example, some mice still are in the phase of compensation stage whereas other mice already show an early sign of heart failure.

PHDs are oxygen sensors, which use molecular O2 as a substrate to hydroxylate proline residues of HIF-α. Deficiency of PHD2 results in stabilization and accumulation of HIF-α, and formation of HIF-α/HIF-β heterodimer, which consequently activates expression of a number of HIF target genes that regulating angiogenesis, inflammation and metabolism^11,13,37^. The cardiac remodeling phenotype in *Egln1*^*Tie2Cre*^ mice is ascribed to activation of HIF-2α but not HIF-1α, as *Hif2a* deletion in EC protects from *Egln1*-deficiency induced cardiac hypertrophy and fibrosis, whereas *Hif1a* deletion in *Egln1*^*Tie2Cre*^ mice does not show any protection but increases cardiac hypertrophy. Consistently, previous studies show that endothelial *Hif1a* deletion in TAC-challenged mice induce myocardial hypertrophy and fibrosis, and rapid decompensation^38^. Although both HIF-1α and HIF-2α express in ECs and share similar DNA binding site, endothelial HIF-1α and HIF-2α plays quite distinct roles in cardiac homeostasis. We speculate that it might be due to the distinct sets of gene activated by HIF-1α and HIF-2α in ECs.

Increased vascular mass and EC proliferation are evident in the LV of *Egln1*^*Tie2Cre*^ mice, demonstrating that PHD2 deficiency in cardiac vascular ECs induces EC proliferation and angiogenesis in mice. Our finding is consistent with previous studies that silencing of endogenous PHD2 in ECs enhances hypoxia-induced EC proliferation^39^. Moreover, previous studies have shown that induction of myocardial angiogenesis promotes cardiomyocyte growth and cardiac hypertrophy^5^. Thus, it is possible that PHD2 deficiency in ECs induces angiogenesis which promotes cardiomyocyte hypertrophy leading to LV hypertrophy. Rarefaction of cardiac microvasculature is associated with pathological hypertrophy^40^. We in fact observed a marked increase of capillary density in *Egln1*^*Tie2Cre*^ hearts, which may help to explain normal cardiac function including fractional shortening and ejection fraction in *Egln1*^*Tie2Cre*^ hearts

The heart is highly organized and consists of multiple cell types including cardiomyocytes, ECs and fibroblasts. Cell-cell communication in the heart is important for cardiac development and adaptation to stress such as pressure overload. It is well documented that soluble factors secreted by ECs maintain tissue homeostasis in different physiological and pathological microenvironment including cancer and bone marrow niche^4,7,41^. To date, a growing number of cardio-active factors derived from ECs have been shown to regulate cardiac angiogenesis contributing to cardiac hypertrophy and/or dysfunction, including Nitric oxide (NO), Neuregulin-1, basic fibroblast growth factor (FGF2), Platelet-derived growth factor (PDGF), Vascular endothelial growth factor (VEGF)^8,41^. Other endothelium-derived factors such as Endothelin-1, NO, neuregulin, *Bmp4, Fgf23, Lgals3, Lcn2, Spp1, Tgfb1*, can directly promote cardiac hypertrophy and fibrosis^42–46^. Our data suggest that endothelial-derived factors mediate EC-myocytes crosstalk to induce cardiac hypertrophy and fibrosis through excessive angiogenesis and/or paracrine effects.

Pathological cardiac hypertrophy worsens clinical outcomes and progresses to heart failure and death. Modulation of abnormal cardiac growth is becoming a potential approach for preventing and treating heart failure in patients^2^. Our study demonstrates that pharmacological inhibition of HIF-2α reduces cardiac hypertrophy in *Egln1*^*Tie2Cre*^ mice, which provides strong evidence that targeting PHD2/HIF-2α signaling is a promising strategy for patients with pathological cardiac hypertrophy. The HIF-2α translation inhibitor C76 used in this study selectively HIF-2α translation by enhancing the binding of iron-regulatory protein 1 to the 5’-untranslated region of *HIF2A* mRNA without affecting HIF-1α expression^31,35^. Inhibition of HIF-1α might be detrimental based on previous and our findings that endothelial HIF-1α is protective in term of cardiac hypertrophy^38^. Moreover, HIF-2α-selective antagonists, which target HIF-2α and HIF-β heterodimerization, are under investigation in renal cancer patients in clinical trials^47–49^. Further studies will be warranted to study this class of HIF-2α inhibitors for the treatment of pathological cardiac hypertrophy and heart failure.

In conclusion, our findings demonstrate for the first time that endothelial PHD2 deficiency in mice induces spontaneous cardiac hypertrophy and fibrosis via HIF-2α activation but not HIF-1α. PHD2 expression was markedly decreased in cardiovascular ECs of patients with cardiomyopathy, validating the clinical relevance of our findings in mice. Thus, selective targeting the abnormality of PHD2/HIF-2α signaling is a potential therapeutic strategy to treat patients with pathological cardiac hypertrophy and fibrosis.

## Author Contributions

Z.D. and Y.Y.Z. conceived the experiments and interpreted the data. Z.D., J.C., B.L., D.Y., C.G., M.M.Z., X.Z., designed and performed experiments, and Z.D., A.F., Y.W. and Y.Y.Z. analyzed the data. Z.D. wrote the manuscript. T.W., Y.Y.Z., revised the manuscript.

---

**Supplemental Table 1.**
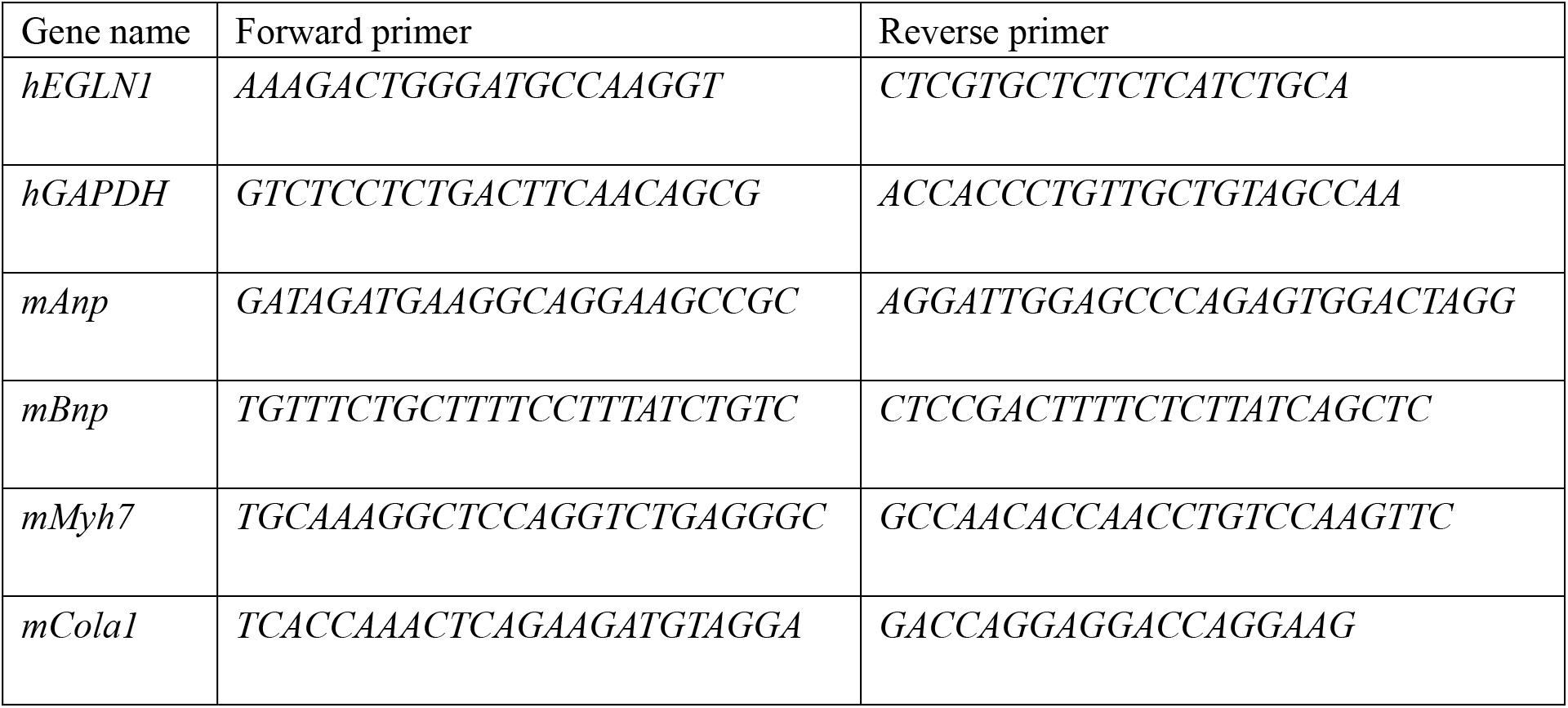
QRT-PCR primers.

**Supplemental Figure 1.**
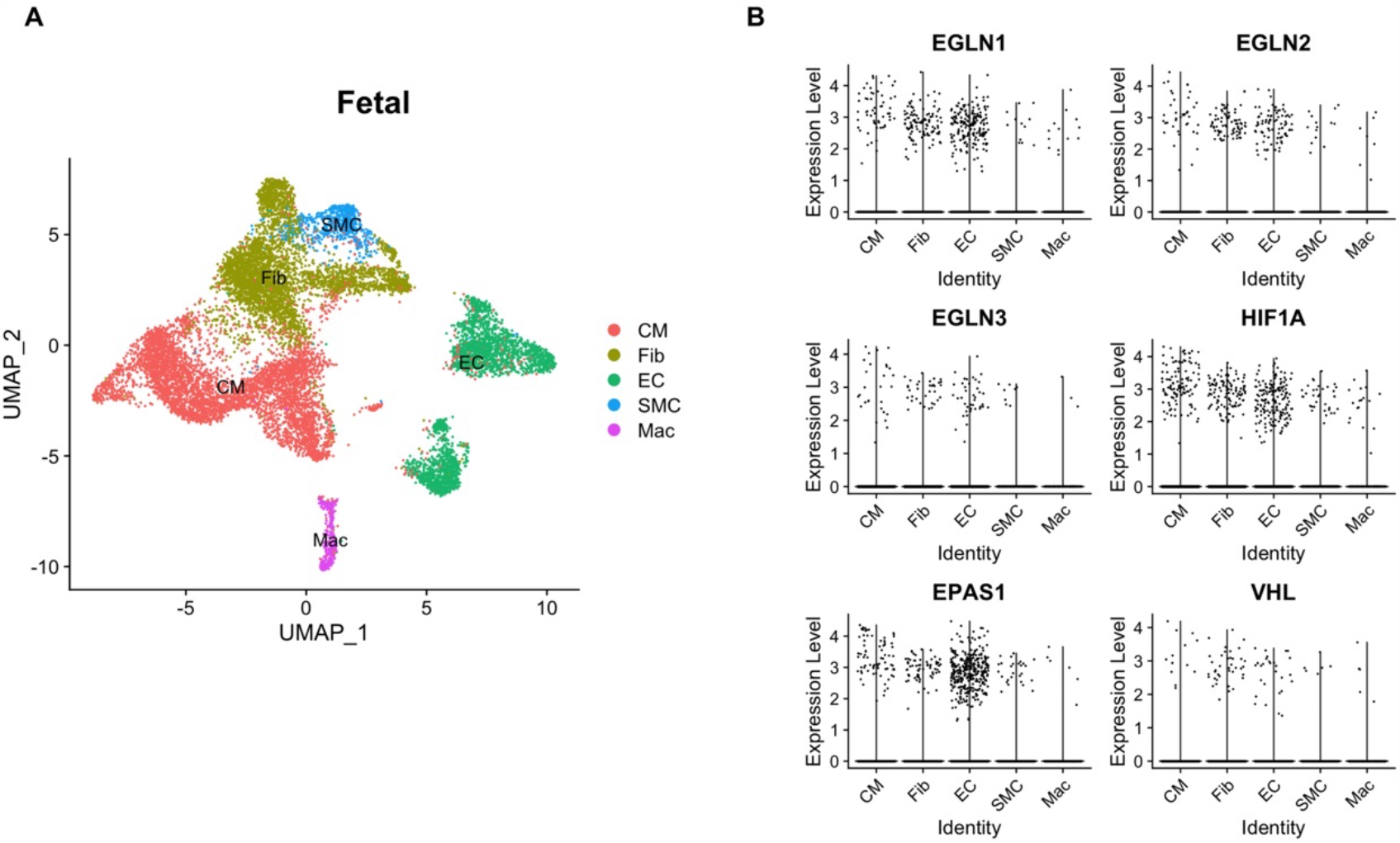
Single-cell RNA sequencing analysis of human fetal hearts. (**A**) A UMAP plot showing the major cardiac cell types including ECs, cardiomyocytes (CM), fibroblasts (Fib), smooth muscle cells (SMC) and macrophages (Mac). (**B**) A Violin plot showing the expression levels of PHD2/HIF signaling genes in cardiac cells.

**Supplemental Figure 2.**
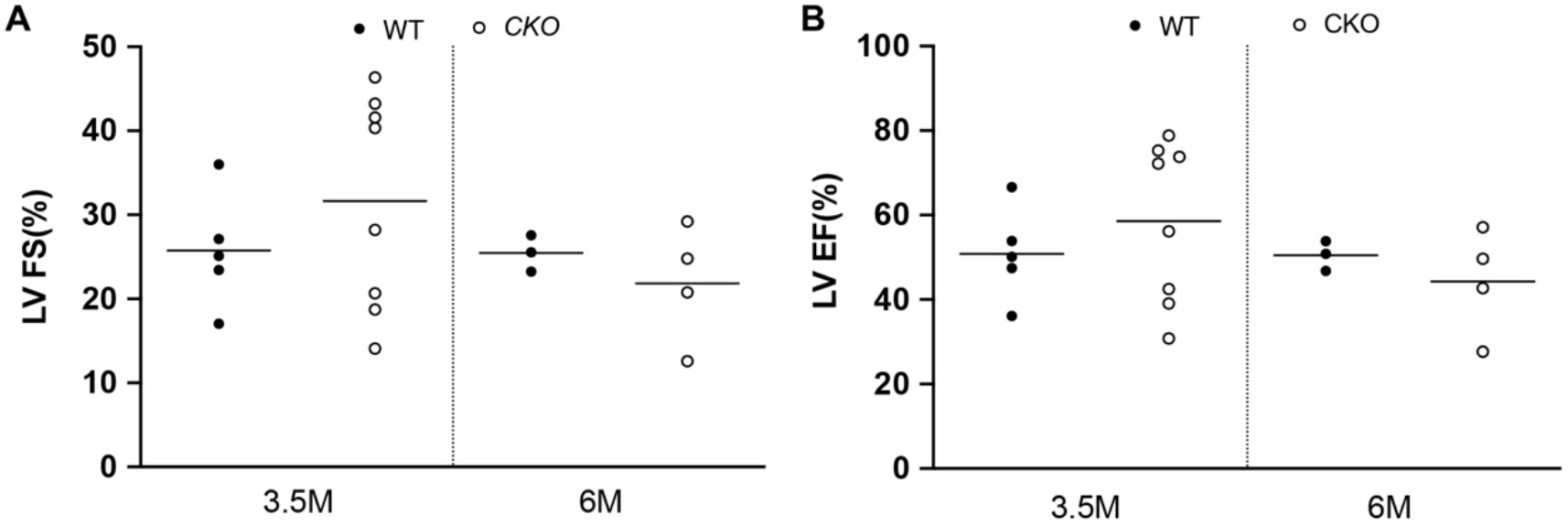
Constitutive loss of endothelial *Egln1* in mice does not affect cardiac function. Echocardiography measurement showing similar LV fraction shorting (FS%) and ejection fraction (EF%) in adult *Egln1*^*Tie2Cre*^ mice compared to WT mice.

**Supplemental Figure 3.**
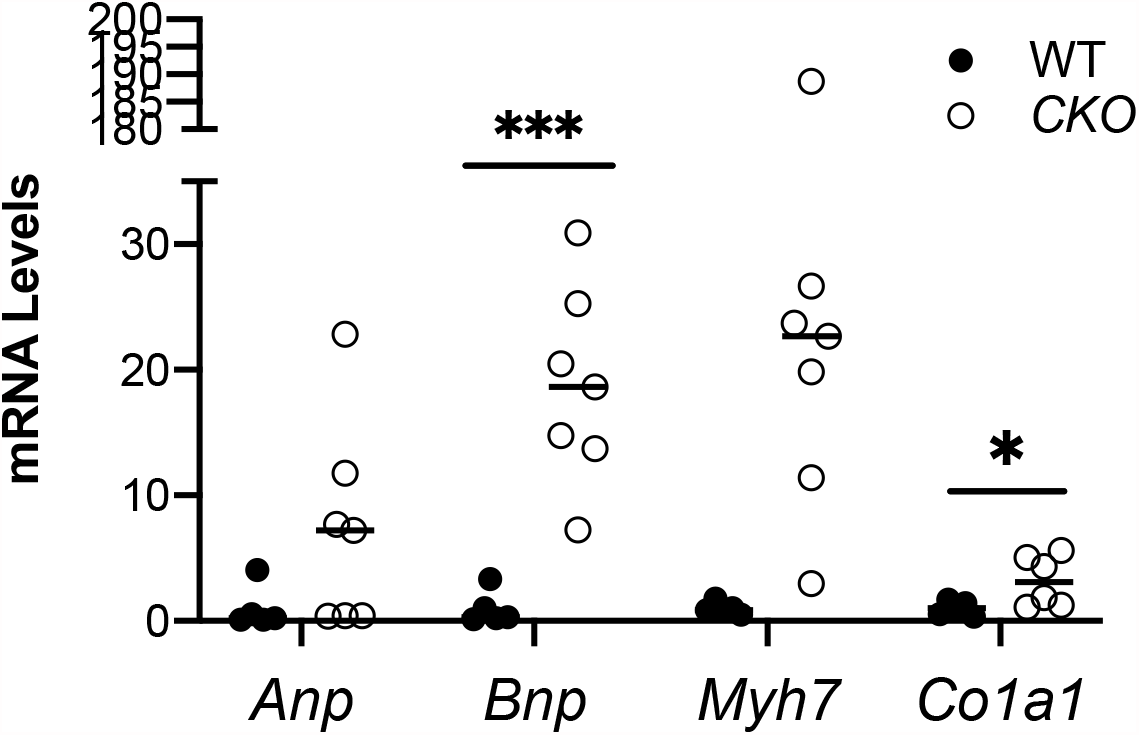
Upregulation of fetal gene including *Bnp* and fibrotic gene *Col1a* in the LV of *Egln1*^*Tie2Cre*^ mice. * p <0.05, ** p <0.01 (Student’s *t* test).

**Supplemental Figure 4.**
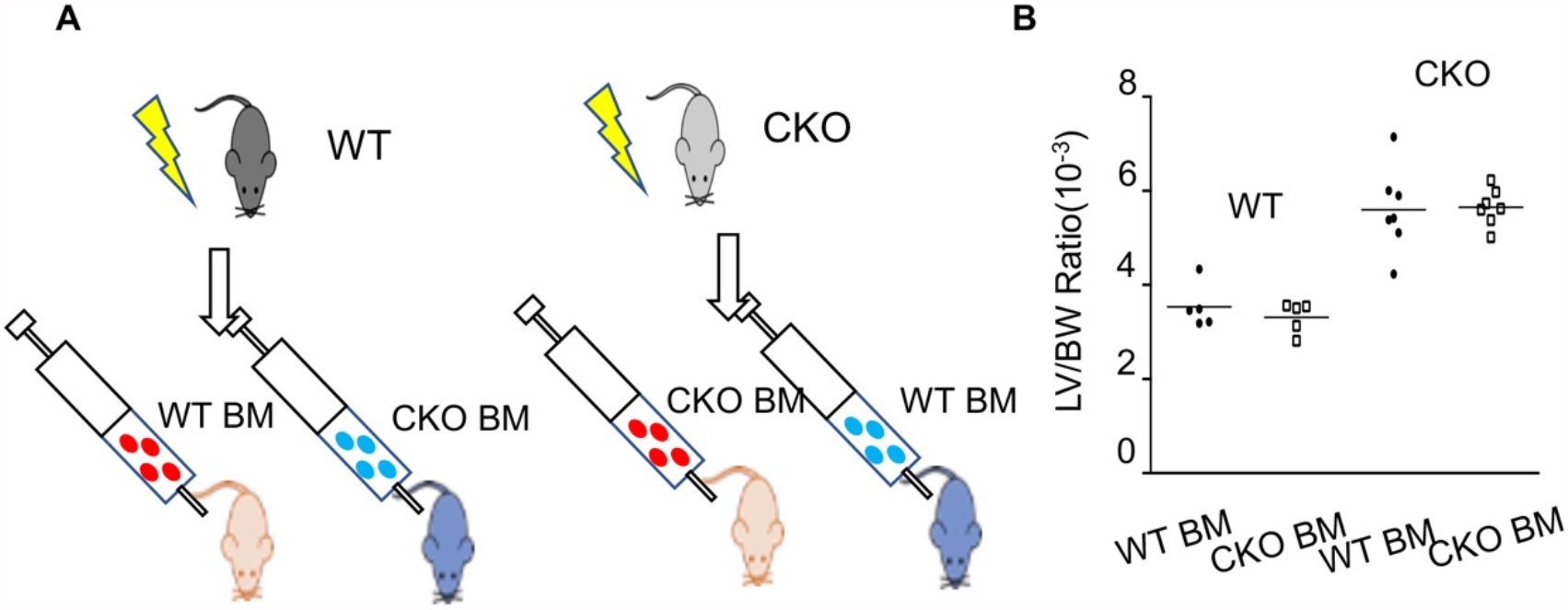
*Egln1* deficiency in bone marrow cells does not contribute to cardiac hypertrophy. (**A**) A diagram showing the strategy of bone marrow cell transplantation. Lethally gamma-irradiated WT were transplanted with bone marrow (BM) cells freshly isolated from WT or *Egln1*^*Tie2Cre*^ (CKO) mice. Similarly, irradiated CKO mice were reconstituted with WT or CKO bone marrow cells. (**B**) Cardiac dissection showed that *Egln1-*deficient bone marrow cells did not contribute to cardiac hypertrophy seen in *Egln1*^*Tie2Cre*^ (CKO) mice.

## Reference

1. Nakamura M, Sadoshima J. Mechanisms of physiological and pathological cardiac hypertrophy. Nat Rev Cardiol. 2018;15:387–407.

2. Frey N, Olson EN. Cardiac Hypertrophy: The Good, the Bad, and the Ugly. Annu Rev Physiol. 2003;65:45–79.

3. Gogiraju R, Schroeter MR, Bochenek ML, Hubert A, Münzel T, Hasenfuss G, Schäfer K. Endothelial deletion of protein tyrosine phosphatase-1B protects against pressure overload-induced heart failure in mice. Cardiovasc Res. 2016;111:204–216.

4. Oka T, Akazawa H, Naito AT, Komuro I. Angiogenesis and Cardiac Hypertrophy. Circ Res. 2014;114:565–571.

5. Tirziu D, Chorianopoulos E, Moodie KL, Palac RT, Zhuang ZW, Tjwa M, Roncal C, Eriksson U, Fu Q, Elfenbein A, Hall AE, Carmeliet P, Moons L, Simons M. Myocardial hypertrophy in the absence of external stimuli is induced by angiogenesis in mice. J Clin Invest. 2007;117:3188–97.

6. Jaba IM, Zhuang ZW, Li N, Jiang Y, Martin KA, Sinusas AJ, Papademetris X, Simons M, Sessa WC, Young LH, Tirziu D. No triggers rgs4 degradation to coordinate angiogenesis and cardiomyocyte growth. J Clin Invest. 2013;123:1718–1731.

7. Shiojima I. Disruption of coordinated cardiac hypertrophy and angiogenesis contributes to the transition to heart failure. J Clin Invest. 2005;115:2108–2118.

8. Talman V, Kivelä R. Cardiomyocyte—Endothelial Cell Interactions in Cardiac Remodeling and Regeneration. Front Cardiovasc Med. 2018;5:1–8.

9. Bassat E, Mutlak YE, Genzelinakh A, Shadrin IY, Baruch Umansky K, Yifa O, Kain D, Rajchman D, Leach J, Riabov Bassat D, Udi Y, Sarig R, Sagi I, Martin JF, Bursac N, Cohen S, Tzahor E. The extracellular matrix protein agrin promotes heart regeneration in mice. Nature. 2017;547:179–184.

10. Epstein ACR, Gleadle JM, McNeill LA, Hewitson KS, O’Rourke J, Mole DR, Mukherji M, Metzen E, Wilson MI, Dhanda A, Tian Y-M, Masson N, Hamilton DL, Jaakkola P, Barstead R, Hodgkin J, Maxwell PH, Pugh CW, Schofield CJ, Ratcliffe PJ. C. elegans EGL-9 and Mammalian Homologs Define a Family of Dioxygenases that Regulate HIF by Prolyl Hydroxylation. Cell. 2001;107:43– 54.

11. Semenza GL. Hypoxia-Inducible Factors in Physiology and Medicine. Cell. 2012;148:399–408.

12. Ivan M, Kondo K, Yang H, Kim W, Valiando J, Ohh M, Salic A, Asara JM, Lane WS, Kaelin WG. HIFalpha Targeted for VHL-Mediated Destruction by Proline Hydroxylation: Implications for O2 Sensing. Science (80-). 2001;292:464–468.

13. Majmundar AJ, Wong WJ, Simon MC. Hypoxia-Inducible Factors and the Response to Hypoxic Stress. Mol Cell. 2010;40:294–309.

14. Lee KE, Simon MC. SnapShot: Hypoxia-Inducible Factors. Cell. 2015;163:1288.e1-1288.e1.

15. Wiesener MS, Jürgensen JS, Rosenberger C, Scholze C, Hörstrup JH, Warnecke C, Mandriota S, Bechmann I, Frei UA, Pugh CW, Ratcliffe PJ, Bachmann S, Maxwell PH, Eckardt K. Widespread, hypoxia-inducible expression of HIF-2α in distinct cell populations of different organs. FASEB J. 2003;17:271–273.

16. Dai Z, Li M, Wharton J, Zhu MM, Zhao Y-Y. Prolyl-4 Hydroxylase 2 (PHD2) Deficiency in Endothelial Cells and Hematopoietic Cells Induces Obliterative Vascular Remodeling and Severe Pulmonary Arterial Hypertension in Mice and Humans Through Hypoxia-Inducible Factor-2α. Circulation. 2016;133:2447–58.

17. Huang X, Zhang X, Zhao DX, Yin J, Hu G, Evans CE, Zhao YY. Endothelial Hypoxia-Inducible Factor-1α Is Required for Vascular Repair and Resolution of Inflammatory Lung Injury through Forkhead Box Protein M1. Am J Pathol. 2019;189:1664–1679.

18. Minamishima YA, Moslehi J, Bardeesy N, Cullen D, Bronson RT, Kaelin WG. Somatic inactivation of the PHD2 prolyl hydroxylase causes polycythemia and congestive heart failure. Blood. 2008;111:3236–3244.

19. Hölscher M, Silter M, Krull S, Von Ahlen M, Hesse A, Schwartz P, Wielockx B, Breier G, Katschinski DM, Zieseniss A. Cardiomyocyte-specific prolyl-4-hydroxylase domain 2 knock out protects from acute myocardial ischemic injury. J Biol Chem. 2011;286:11185–11194.

20. Fan Q, Mao H, Angelini A, Coarfa C, Robertson MJ, Lagor WR, Wehrens XHT, Martin JF, Pi X, Xie L. Depletion of Endothelial Prolyl Hydroxylase Domain Protein 2 and 3 Promotes Cardiomyocyte Proliferation and Prevents Ventricular Failure Induced by Myocardial Infarction. Circulation. 2019;140:440–442.

21. Gao C, Ren SV, Yu J, Baal U, Thai D, Lu J, Zeng C, Yan H, Wang Y. Glucagon Receptor Antagonism Ameliorates Progression of Heart Failure. JACC Basic to Transl Sci. 2019;4:161–172.

22. Wang Z, Zhang XJ, Ji YX, Zhang P, Deng KQ, Gong J, Ren S, Wang X, Chen I, Wang H, Gao C, Yokota T, Ang YS, Li S, Cass A, Vondriska TM, Li G, Deb A, Srivastava D, Yang HT, Xiao X, Li H, Wang Y. The long noncoding RNA Chaer defines an epigenetic checkpoint in cardiac hypertrophy. Nat Med. 2016;22:1131–1139.

23. Tran KA, Zhang X, Predescu D, Huang X, Machado RF, Göthert JR, Malik AB, Valyi-Nagy T, Zhao Y-Y. Endothelial β-Catenin Signaling Is Required for Maintaining Adult Blood–Brain Barrier Integrity and Central Nervous System Homeostasis. Circulation. 2016;133:177–186.

24. Han X, Zhou Z, Fei L, Sun H, Wang R, Chen Y, Chen H, Wang J, Tang H, Ge W, Zhou Y, Ye F, Jiang M, Wu J, Xiao Y, Jia X, Zhang T, Ma X, Zhang Q, Bai X, Lai S, Yu C, Zhu L, Lin R, Gao Y, Wang M, Wu Y, Zhang J, Zhan R, Zhu S, Hu H, Wang C, Chen M, Huang H, Liang T, Chen J, Wang W, Zhang D, Guo G. Construction of a human cell landscape at single-cell level. Nature. 2020;581:303– 309.

25. Stuart T, Butler A, Hoffman P, Hafemeister C, Papalexi E, Mauck WM, Hao Y, Stoeckius M, Smibert P, Satija R. Comprehensive Integration of Single-Cell Data. Cell. 2019;177:1888–1902.e21.

26. Chen J, Sathiyamoorthy K, Zhang X, Schaller S, Perez White BE, Jardetzky TS, Longnecker R. Ephrin receptor A2 is a functional entry receptor for Epstein-Barr virus. Nat Microbiol. 2018;3:172– 180.

27. Göthert JR, Gustin SE, Van Eekelen JAM, Schmidt U, Hall MA, Jane SM, Green AR, Göttgens B, Izon DJ, Begley CG. Genetically tagging endothelial cells in vivo: Bone marrow-derived cells do not contribute to tumor endothelium. Blood. 2004;104:1769–1777.

28. Nussbaum C, Bannenberg S, Keul P, Gräler MH, Gonçalves-De-Albuquerque CF, Korhonen H, Von Wnuck Lipinski K, Heusch G, De Castro Faria Neto HC, Rohwedder I, Göthert JR, Prasad VP, Haufe G, Lange-Sperandio B, Offermanns S, Sperandio M, Levkau B. Sphingosine-1-phosphate receptor 3 promotes leukocyte rolling by mobilizing endothelial P-selectin. Nat Commun. 2015;6.

29. Weis SM, Lim ST, Lutu-Fuga KM, Barnes LA, Chen XL, Göthert JR, Shen TL, Guan JL, Schlaepfer DD, Cheresh DA. Compensatory role for Pyk2 during angiogenesis in adult mice lacking endothelial cell FAK. J Cell Biol. 2008;181:43–50.

30. Cheng KT, Xiong S, Ye Z, Hong Z, Di A, Tsang KM, Gao X, An S, Mittal M, Vogel SM, Miao EA, Rehman J, Malik AB. Caspase-11-mediated endothelial pyroptosis underlies endotoxemia-induced lung injury. J Clin Invest. 2017;127:4124–4135.

31. Dai Z, Zhu MM, Peng Y, Machireddy N, Evans CE, Machado R, Zhang X, Zhao Y-Y. Therapeutic Targeting of Vascular Remodeling and Right Heart Failure in Pulmonary Arterial Hypertension with a HIF-2α Inhibitor. Am J Respir Crit Care Med. 2018;198:1423–1434.

32. Dai Z, Zhu MM, Peng Y, Jin H, Machireddy N, Qian Z, Zhang X, Zhao Y-Y. Endothelial and Smooth Muscle Cell Interaction via FoxM1 Signaling Mediates Vascular Remodeling and Pulmonary Hypertension. Am J Respir Crit Care Med. 2018;198:788–802.

33. Wu S, Nishiyama N, Kano MR, Morishita Y, Miyazono K, Itaka K, Chung U Il, Kataoka K. Enhancement of angiogenesis through stabilization of hypoxia-inducible factor-1 by silencing prolyl hydroxylase domain-2 gene. Mol Ther. 2008;16:1227–1234.

34. Chan DA, Kawahara TLA, Sutphin PD, Chang HY, Chi JT, Giaccia AJ. Tumor Vasculature Is Regulated by PHD2-Mediated Angiogenesis and Bone Marrow-Derived Cell Recruitment. Cancer Cell. 2009;15:527–538.

35. Zimmer M, Ebert BL, Neil C, Brenner K, Papaioannou I, Melas A, Tolliday N, Lamb J, Pantopoulos K, Golub T, Iliopoulos O. Small-Molecule Inhibitors of HIF-2a Translation Link Its 5′UTR Iron-Responsive Element to Oxygen Sensing. Mol Cell. 2008;32:838–848.

36. Frey N, Katus HA, Olson EN, Hill JA. Hypertrophy of the Heart: A New Therapeutic Target? Circulation. 2004;109:1580–1589.

37. Bishop T, Ratcliffe PJ. HIF Hydroxylase Pathways in Cardiovascular Physiology and Medicine. Circ Res. 2015;117:65–79.

38. Wei H, Bedja D, Koitabashi N, Xing D, Chen J, Fox-Talbot K, Rouf R, Chen S, Steenbergen C, Harmon JW, Dietz HC, Gabrielson KL, Kass DA, Semenza GL. Endothelial expression of hypoxia-inducible factor 1 protects the murine heart and aorta from pressure overload by suppression of TGF-signaling. Proc Natl Acad Sci. 2012;109:E841–E850.

39. Takeda K, Fong GH. Prolyl hydroxylase domain 2 protein suppresses hypoxia-induced endothelial cell proliferation. Hypertension. 2007;49:178–184.

40. Mohammed SF, Hussain S, Mirzoyev SA, Edwards WD, Maleszewski JJ, Redfield MM. Coronary microvascular rarefaction and myocardial fibrosis in heart failure with preserved ejection fraction. Circulation. 2015;131:550–559.

41. Gogiraju R, Bochenek ML, Schäfer K. Angiogenic Endothelial Cell Signaling in Cardiac Hypertrophy and Heart Failure. Front Cardiovasc Med. 2019;6.

42. Grabner A, Schramm K, Silswal N, Hendrix M, Yanucil C, Czaya B, Singh S, Wolf M, Hermann S, Stypmann J, Marco GS DI, Brand M, Wacker MJ, Faul C. FGF23/FGFR4-mediated left ventricular hypertrophy is reversible. Sci Rep. 2017;7:1–12.

43. Sun B, Huo R, Sheng Y, Li Y, Xie X, Chen C, Liu H Bin, Li N, Li CB, Guo WT, Zhu JX, Yang BF, Dong DL. Bone morphogenetic protein-4 mediates cardiac hypertrophy, apoptosis, and fibrosis in experimentally pathological cardiac hypertrophy. Hypertension. 2013;61:352–360.

44. de Boer RA, Edelmann F, Cohen-Solal A, Mamas MA, Maisel A, Pieske B. Galectin-3 in heart failure with preserved ejection fraction. Eur J Heart Fail. 2013;15:1095–1101.

45. Khalil H, Kanisicak O, Prasad V, Correll RN, Fu X, Schips T, Vagnozzi RJ, Liu R, Huynh T, Lee S-J, Karch J, Molkentin JD. Fibroblast-specific TGF-β–Smad2/3 signaling underlies cardiac fibrosis. J Clin Invest. 2017;127:3770–3783.

46. López B, González A, Lindner D, Westermann D, Ravassa S, Beaumont J, Gallego I, Zudaire A, Brugnolaro C, Querejeta R, Larman M, Tschöpe C, Díez J. Osteopontin-mediated myocardial fibrosis in heart failure: A role for lysyl oxidase? Cardiovasc Res. 2013;99:111–120.

47. Cho H, Du X, Rizzi JP, Liberzon E, Chakraborty AA, Gao W, Carvo I, Signoretti S, Bruick RK, Josey JA, Wallace EM, Kaelin WG. On-target efficacy of a HIF-2α antagonist in preclinical kidney cancer models. Nature. 2016;539:107–111.

48. Chen W, Hill H, Christie A, Kim MS, Holloman E, Pavia-Jimenez A, Homayoun F, Ma Y, Patel N, Yell P, Hao G, Yousuf Q, Joyce A, Pedrosa I, Geiger H, Zhang H, Chang J, Gardner KH, Bruick RK, Reeves C, Hwang TH, Courtney K, Frenkel E, Sun X, Zojwalla N, Wong T, Rizzi JP, Wallace EM, Josey JA, Xie Y, Xie XJ, Kapur P, McKay RM, Brugarolas J. Targeting renal cell carcinoma with a HIF-2 antagonist. Nature. 2016;539:112–117.

49. Courtney KD, Infante JR, Lam ET, Figlin RA, Rini BI, Brugarolas J, Zojwalla NJ, Lowe AM, Wang K, Wallace EM, Josey JA, Choueiri TK. Phase I Dose-Escalation Trial of PT2385, a First-in-Class Hypoxia-Inducible Factor-2α Antagonist in Patients With Previously Treated Advanced Clear Cell Renal Cell Carcinoma. J Clin Oncol. 2018;36:867–874.

